# Beyond the Final Layer: Evaluating Single-Cell Foundation Model Representations for Molecular Prediction

**DOI:** 10.64898/2026.10.05.756880

**Authors:** Ming-Ching Crystal Wen, YiWei Lyu, Andrew Robbins

## Abstract

Single-cell foundation models (scFMs) are commonly benchmarked using final-layer embeddings, although useful depth may depend on the task. We evaluate Geneformer, scBERT, scGPT, and scFoundation as frozen feature extractors for cell-type and perturbation-response prediction using layer-wise probes, leave-one-dataset-out (LODO) selection, readout controls, representation diagnostics, centered kernel alignment (CKA), and retrieval/collapse analyses. Intermediate layers significantly improve cell-type accuracy for scGPT and scBERT in several settings, while Geneformer and scFoundation generally show performance comparable to their final layers. For perturbation prediction, LODO identifies layer 1 as an earliest-on-par candidate across all four models, without establishing universal superiority. Paired scGPT CLS and mean-pooling analyses show that readout affects preferred depth. CKA and representation metrics reveal model- and task-dependent geometric changes, while retrieval and collapse checks support the robustness of perturbation findings. We recommend reporting extraction depth, readout, selection protocol, and task-specific diagnostics. A validated non-final layer may also reduce computation, although efficiency was not directly benchmarked.

## 1 Introduction

Modern molecular-science workflows increasingly compose learned representations with downstream predictors, retrieval modules, and decision procedures. In single-cell biology, single-cell foundation models (scFMs) now serve this role for cell annotation, perturbation analysis, and downstream biological prediction [1–4]. A common benchmarking shortcut is to extract one embedding per model, usually from the final layer. That shortcut implicitly assumes that the last hidden representation is the most useful representation for molecular and cellular prediction. We test this assumption directly.

We evaluate four influential scFMs: Geneformer, scBERT, scGPT, and scFoundation [1–4]. Instead of fixing one extraction layer, we probe all available layers on two tasks aligned with molecular-science benchmarking: cell type prediction and prediction of perturbation-induced gene expression shifts. This matters because a benchmark, or a larger agentic workflow that consumes its embeddings, can rank models or draw biological conclusions differently depending on which representation layer is used for the task being measured.

Our contribution is an evaluation pipeline and empirical case study showing that extraction depth changes both absolute performance and practical layer-selection recommendations. The key point is not that one intermediate layer should replace the final layer everywhere, but that final-layer-only benchmarking is an under-specified evaluation protocol, especially given evidence that scFM evaluation is sensitive to task, readout, and benchmark design [5–10]. We therefore combine layer-wise probing with LODO layer selection, paired scGPT CLS-versus-mean readout controls, cross-dataset cell-type transfer, and perturbation retrieval/collapse diagnostics. Together, these analyses distinguish statistically supported intermediate-layer gains, practically on-par early layers, and settings where the final layer remains the conservative choice.

## 2 Methods

### Layer-wise probing

All models are evaluated as frozen feature extractors. For cell type prediction, we use four datasets (CZI human meninges, PBMC3K, Tabula Sapiens testis, and Zheng68k) [11–16], remove rare classes, extract one cell embedding per layer using each model’s native readout, and train multinomial logistic-regression probes with 5-fold cross-validation. We report both accuracy and macro F1 to capture overall predictive performance while giving minority cell types equal weight in the evaluation. For perturbation prediction, we use three perturbation datasets, including two CRISPR-based genetic datasets and the sci-Plex chemical transcriptomics dataset [17–19] spanning genetic and chemical perturbations, average cell embeddings within each perturbation, predict mean expression deltas relative to control using multi-output ridge regression, and report Pearson correlation coefficient (PCC)-delta and mean square error (MSE)-delta.

### Layer selection and statistical tests

For cell type LODO selection, each target dataset is excluded from layer selection. The selected layer is chosen using the remaining datasets after averaging folds within each dataset, with ties resolved toward earlier layers. We then compare the selected layer with the final layer using paired cell-level out-of-fold predictions. Paired accuracy differences are tested with exact McNemar tests over 16 model–dataset comparisons and jointly corrected using BH. Macro-F1 differences are reported as effect sizes with class-stratified bootstrap intervals.

For perturbation LODO selection, each perturbation dataset is held out, and layers are selected using the remaining datasets with prespecified on-par margins: PCC-delta no more than 0.01 below final-layer performance and MSE-delta no more than 5% above final-layer error. We also compute top-1 retrieval and mean reciprocal rank by matching each predicted held-out perturbation delta to observed perturbation deltas. Collapse warnings are triggered by predicted-to-observed variance ratio below 0.10, predicted-to-observed pairwise-distance ratio below 0.10, or fewer than 25% unique Pearson top-1 assignments.

### Readout and transfer controls

Because scGPT natively uses a learned CLS token while some other models are commonly summarized by mean pooling, we repeat scGPT extraction with paired CLS and non-learned mean-pooling readouts using identical cells, preprocessing, folds, probes, and metrics. To test whether cell type prediction conclusions persist beyond within-dataset cross-validation, we also run bidirectional PBMC3K–Zheng68k label transfer after harmonizing seven shared cell types. We use this pair because both are independent PBMC datasets with broad shared immune-cell annotations, enabling a controlled cross-dataset transfer test without the tissue mismatch of Testis or the broader compositional heterogeneity of CZI. Layers are selected using source-only cross-validation before fitting the final source predictor and evaluating on the target dataset.

### Representation diagnostics

We compute layer-wise entropy, effective rank, intrinsic dimension, and CKA (Appendix Figures 1–3). These diagnostics are not universal selectors; they indicate whether useful task representations occur in compressed, expanded, stable, or strongly reorganized regimes, following broader layer-wise analyses of Transformer representations [20–24].

## 3 Results

### 3.1 Cell type prediction shows on-par or stronger non-final layers

Cell type prediction shows clear architecture-dependent depth profiles (Figure 1a). Geneformer is strong from early-to-middle layers onward, scBERT is comparatively depth-stable and can appear late/final-layer favored in the averaged descriptive curve, scFoundation improves toward later layers, and scGPT has the clearest non-final optimum. Across models, an important baseline observation is that non-final layers often reach performance comparable to the final layer even when statistically stronger intermediate-layer performance is not established. Thus, Figure 1a should not be read as claiming that every model has an average non-final peak. Rather, it motivates the stricter selection-controlled analyses in Table 1. In the global best-versus-final analysis, scGPT shows significant non-final advantages for both accuracy (Δ = 0.026, *q* = 4.21 *×* 10^*−*10^) and macro F1 (Δ = 0.106, *q* = 7.22 *×* 10^*−*11^).

**Figure 1.**
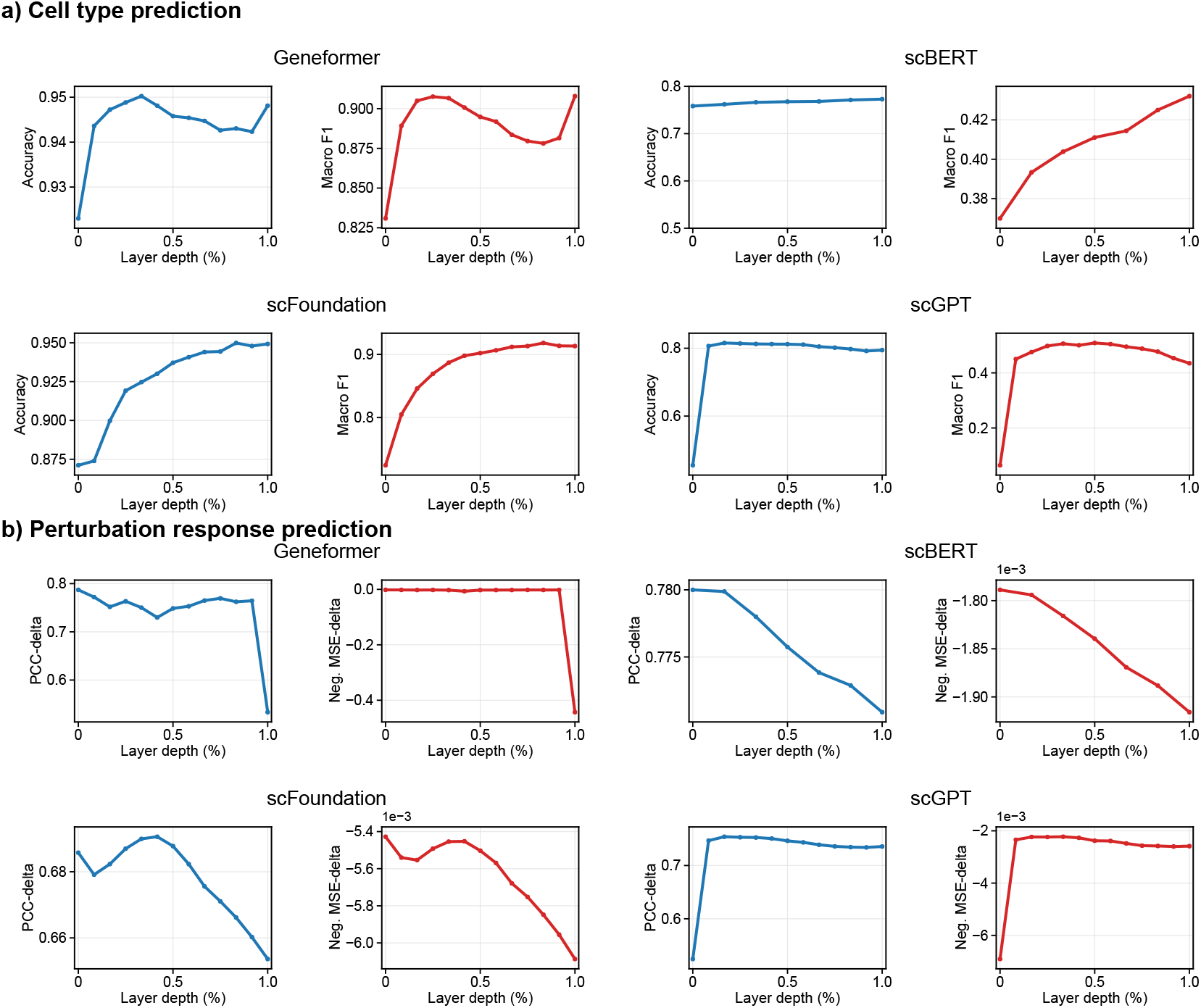
Descriptive layer-wise downstream performance from saved activations. Panel a shows cell-type prediction averaged across four datasets; only the scBERT y-axis is widened to avoid visually exaggerating its small depth effect. Panel b shows perturbation-response prediction averaged across three datasets using PCC-delta and *negative* MSE-delta (MSE), so higher values indicate better performance for both metrics. These target-dataset profiles complement the selection-controlled LODO and global best-versus-final tests summarized in Table 1.

**Table 1.** Evidence map for layer-wise conclusions that summarizes selection-controlled LODO comparisons, corrected best-versus-final tests, and added controls.

| Analysis | Geneformer | scBERT | scGPT | scFoundation |
| --- | --- | --- | --- | --- |
| Cell type LODO | final conserva-<br>tive | non-final + | layer 2 ++ | final conserva-<br>tive |
| Cell type best-vs-final | comparable | modest + | strong ++ | comparable |
| Perturbation best-vs-final | PCC-delta ++ | stable | MSE suggestive | MSE-delta ++ |
| Perturbation LODO | layer 1 on-par | layer 1 on-par | readout-<br>dependent | layer 1 on-par |
| Readout control | – | – | CLS/mean differ | – |
| Transfer control | mixed | mixed | layer 2 transfers | direction-<br>dependent |
| Retrieval/collapse | retrieval + | retrieval + | retrieval + | retrieval + |

++ = corrected non-final advantage; + = smaller or descriptive non-final benefit; on-par = earliest layer satisfying the prespecified final-layer margin; stable/comparable = no strong evidence against the final-layer baseline.

LODO selection asks a different question from the descriptive curves: if the target dataset is excluded from layer choice, does the selected layer beat the final layer on paired out-of-fold predictions? Under this criterion, scGPT layer 2 improves accuracy over the final layer by 3.30, 3.45, 0.55, and 2.82 percentage points on CZI, PBMC3K, Testis, and Zheng68k, respectively; macro-F1 gains are 8.04, 6.49, 8.48, and 8.10 points. scBERT selected intermediate layers show smaller paired gains of 1.60, 1.63, 0.28, and 1.27 points across the same datasets, even though the averaged descriptive curve is relatively flat and late-layer competitive. For both scGPT and scBERT, paired accuracy gains remain significant after correction on CZI, PBMC3K, and Zheng68k, but not Tabula Sapiens testis data. Geneformer and scFoundation show comparable performance between selected non-final layers and final layers under this paired LODO analysis.

The paired gains are biologically structured rather than only aggregate-score changes. Comparing cell-level predictions from the LODO-selected and final layers, scGPT rescues more cells than it harms on all four cell-type datasets (net rescued cells: +279, +91, +41, +436 on CZI, PBMC3K, Tabula Sapiens testis, and Zheng68k data). scBERT shows the same direction (+135, +43, +21, +197). Examples include scGPT recall gains for CZI innate lymphoid cells and Zheng68k plasmacytoid dendritic cells, and scBERT recall gains for CZI megakaryocytes, CZI B cells, and PBMC3K CD8 T cells. Thus, the strongest LODO effects are not merely numerical artifacts of majority classes.

### 3.2 Perturbation prediction has dataset- and metric-dependent optima

Perturbation prediction has different depth behavior from cell type prediction (Figure 1b), consistent with recent cautions that perturbation-response benchmarks require strong baselines and careful validation [25]. Geneformer peaks in shallow layers and drops sharply at the final layer. scBERT remains relatively stable. scFoundation peaks in shallow-to-middle layers and then declines. scGPT improves after the input layer, peaks early-to-middle, and then remains comparatively stable. Notably, the significant best-versus-final tests identify Geneformer PCC-delta (*q*_BH_ = 0.004) and scFoundation MSE-delta (*q*_BH_ = 0.003) even after BH correction.

The LODO perturbation analysis does not justify a single universally best layer. Instead, layer 1 is the earliest aggregate layer satisfying the prespecified on-par criteria relative to the final layer. Although some non-final layers show higher PCC-delta, lower MSE-delta, or stronger retrieval performance than the final layer in individual settings, the preferred layer varies across model, dataset, metric, and scGPT readout. We therefore interpret early layers as practical candidate extraction points that still require target-dataset validation.

This is the main contrast with cell type prediction. For perturbation response, LODO selection tests whether a layer chosen without access to the target dataset recovers the same layer that would be preferred by target-dataset search. Our results do not support a universal perturbation layer in that strong sense. Instead, they support a more cautious evaluation protocol: begin with early layers, verify PCC-delta and MSE-delta against the final layer under prespecified margins, and then check retrieval and collapse diagnostics before treating the representation as reliable.

Practically, these results strongly prioritize scGPT layer 2 for cell type prediction validation and suggest checking scBERT layers 1–4 alongside the final layer, while retaining the final layer as a conservative option for Geneformer and scFoundation when intermediate-layer superiority is not established. For perturbation tasks, early extraction should be treated as a candidate workflow: start at layer 1, validate PCC-delta and MSE-delta against the final layer, and report the readout because scGPT CLS favors layer 4 whereas mean pooling favors layers 1–2.

### 3.3 Geometry explains why depth sensitivity differs

CKA and representation diagnostics help explain why layer choice matters more for some models (Appendix Figures 1–3). scBERT has highly homogeneous cross-layer similarity, matching its weak depth dependence. Geneformer changes more smoothly but loses perturbation-relevant information at the final layer. scFoundation shows a clearer early-to-late shift. scGPT has the strongest early-to-middle reorganization, consistent with its pronounced non-final optima.

The paired scGPT readout analysis further separates readout choice from depth effects (Figure 2). CLS and mean-pooling embeddings yield positively correlated layer-performance profiles for cell type prediction (*ρ* = 0.65–0.88), and same-layer CLS/mean CKA is high for perturbation settings (0.86–0.99). However, the exact optimum changes: in perturbation prediction, CLS favors layer 4 across all three datasets, while mean pooling favors layers 1–2. Thus, readout matters, but useful non-final signal is not an artifact of the CLS token alone.

**Figure 2.**
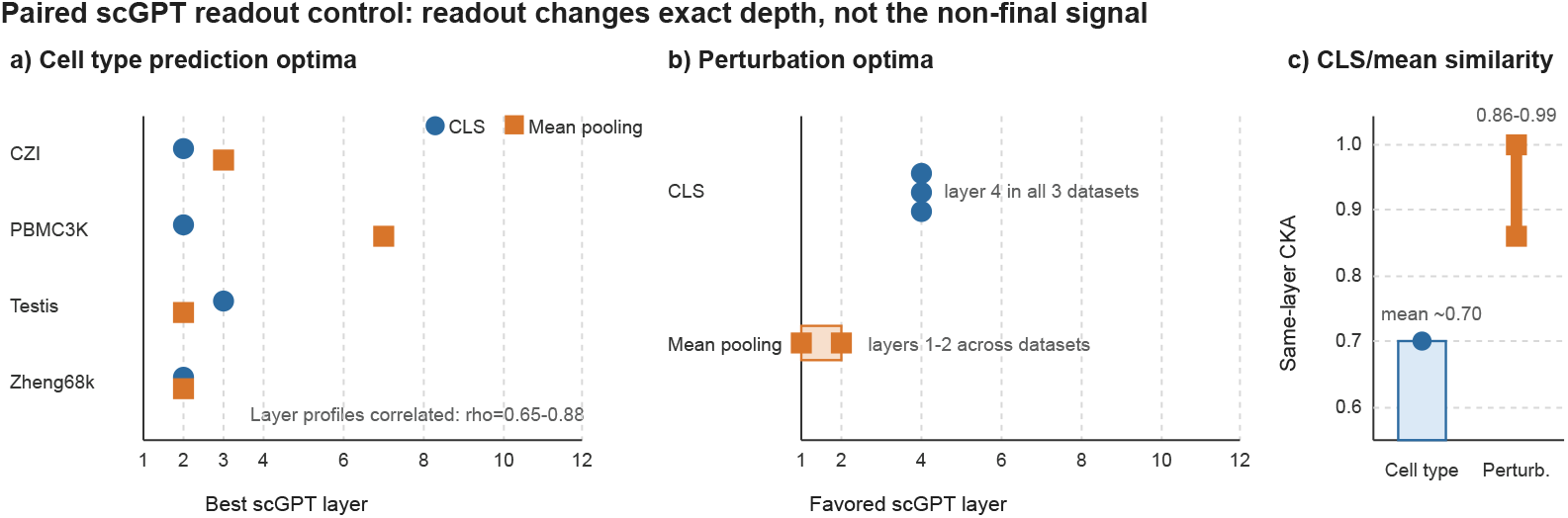
Paired scGPT readout control. CLS and mean-pooling readouts use identical cells, preprocessing, folds, probes, and metrics.

The transfer control gives a complementary view of generalization. In bidirectional PBMC3K– Zheng68k transfer, scGPT layer 2 improves transfer macro F1 in both directions under both CLS and mean pooling. The geometry linked to successful transfer differs by model and readout. For scFoundation, transfer macro F1 is most associated with same-cell-type neighborhood purity, defined as the fraction of nearest neighbors sharing the same cell type label (*ρ* = 0.902, *q*_BH_ *<* 0.001). For scGPT CLS, transfer macro F1 is instead associated with within-cell-type cross-dataset mixing, meaning that cells of the same type from PBMC3K and Zheng68k are interleaved rather than separated by dataset (*ρ* = 0.839, *q*_BH_ = 0.004). Thus, different scFMs can support transfer through different representation geometries.

These geometric results are useful because they show why a single diagnostic is unlikely to solve layer selection. A high CKA-to-final layer can preserve information useful for a final-layer-like readout, but a lower-similarity intermediate layer may better preserve task-relevant variation that later layers compress away. Likewise, higher effective rank or intrinsic dimension can indicate richer variation, but richness alone is not equivalent to biological utility. Thus, the problem is not simply that any single diagnostic may be imperfect; it is that no one diagnostic, readout, or extraction depth should be assumed to work best across all tasks.

## 4 Discussion

The main lesson is methodological: scFM benchmarks should treat extraction depth as an explicit evaluation variable. This matters beyond standalone benchmarking because scFM embeddings may be used inside molecular-science systems for perturbation modeling, retrieval, or experimental prioritization. Final-layer embeddings are a reasonable baseline, but may not be a reliable default. In cell type prediction, LODO selection gives the clearest statistically supported non-final gains for scGPT and scBERT; in perturbation prediction, non-final advantages are more dependent on the dataset and metric.

The controls also narrow the claim appropriately. PBMC3K–Zheng68k transfer shows that scGPT layer 2 can improve cross-dataset cell type prediction transfer, but this remains a cell type prediction generalization setting rather than a comprehensive benchmark over biological objectives. Perturbation retrieval confirms that non-final effects are not simply collapse artifacts, but it also reinforces that perturbation layer selection should be validated for the target dataset and metric.

For benchmark design, final-layer-only reporting is incomplete, much like reporting a single metric. Layer-wise performance, readout, LODO-selected versus target-searched optima, and perturbation retrieval/collapse diagnostics give a more faithful picture of when a pretrained scFM representation is useful. Limitations include four transformer-based scFMs, two objective tasks, frozen single-seed probes, and primarily cell-level rather than donor/batch-grouped inference. Prospective perturbation prediction and direct early-exit savings were not evaluated; our perturbation task decodes responses from already perturbed profiles.

For practitioners, we suggest a lightweight workflow: report the final layer as a baseline, probe a small early/middle/late layer set, validate perturbation settings with prediction and retrieval/collapse metrics, and state the readout. When validated non-final layers are on par with or better than final layers, early extraction can reduce later feed-forward computation and storage for saved activations. This matters for agentic or closed-loop molecular workflows, where downstream tools inherit the benchmark’s extraction choice.

## Code Availability

All code for layer-wise scFM extraction, probing, representation analysis, and reproducibility checks is publicly available at: https://github.com/mccwen/scFM-layer-evaluation.

## Appendix Representation Diagnostics and CKA

### Additional readout, transfer, and biological-rescue analyses

The paired scGPT readout analysis uses identical cells, preprocessing, folds, probes, and metrics for the native CLS state and non-learned mean pooling. CLS and mean-pooling layer-performance profiles for cell type prediction have Spearman correlations of *ρ* = 0.65–0.88. In perturbation prediction, CLS favors layer 4 across all three datasets, whereas mean pooling favors layers 1–2. Same-layer CLS/mean CKA is approximately 0.70 for cell-type datasets and 0.86–0.99 for perturbation datasets. Thus, readout changes the exact recommendation but does not remove useful non-final signal.

In bidirectional PBMC3K–Zheng68k transfer, scGPT layer 2 improves macro F1 in both directions under both CLS and mean pooling. Transfer macro F1 is associated with cell-type purity for scFoundation (*ρ* = 0.902, *q*_BH_ *<* 0.001), within-cell-type cross-dataset mixing for scGPT CLS (*ρ* = 0.839, *q*_BH_ = 0.004), and cell-type purity for scGPT mean pooling (*ρ* = 0.699, *q*_BH_ = 0.049). These results are a narrow transfer control, not a complete integration benchmark.

Cell-level OOF error analysis showed that scGPT intermediate layers had net rescued cells of +279, +91, +41, +436 across CZI, PBMC3K, Testis, and Zheng68k, while scBERT had

+135, +43, +21, +197. Examples included improved recall for CZI innate lymphoid cells and Zheng68k plasmacytoid dendritic cells with scGPT, and CZI megakaryocytes, CZI B cells, and PBMC3K CD8 T cells with scBERT. These structured gains should be interpreted as cell-level evidence, not donor-level replication.

### Selection-controlled supplementary results

Appendix Table 1 gives the compact cell-type LODO summary used in the main text. Accuracy changes are selected-layer minus final-layer percentage points; *q*_BH_ refers to the paired accuracy test. Geneformer and scFoundation had no BH-significant intermediate-layer advantage in the reported comparisons.

**Appendix Table 1.** Cell-type LODO effects. Each layer was selected without using the held-out dataset.

| Model | Held-out data | Layer | Accuracy $\Delta$ | BH result |
| --- | --- | --- | --- | --- |
| scGPT | CZI | 2 | +3.30 | significant |
| scGPT | PBMC3K | 2 | +3.45 | significant |
| scGPT | Testis | 2 | +0.55 | not significant |
| scGPT | Zheng68k | 2 | +2.82 | significant |
| scBERT | CZI | 1/4 | +1.60 | significant |
| scBERT | PBMC3K | 1/4 | +1.63 | significant |
| scBERT | Testis | 1/4 | +0.28 | not significant |
| scBERT | Zheng68k | 1/4 | +1.27 | significant |
| Geneformer | all | dataset-dependent | comparable | none significant |
| scFoundation | all | 10 or final | comparable | none significant |

Appendix Table 2 reports the held-out perturbation LODO ranges across three datasets. Layer 1 was selected for every model as the earliest layer satisfying both prespecified on-par criteria. Positive PCC differences favor layer 1; negative MSE differences indicate lower error for layer 1.

**Appendix Table 2.**
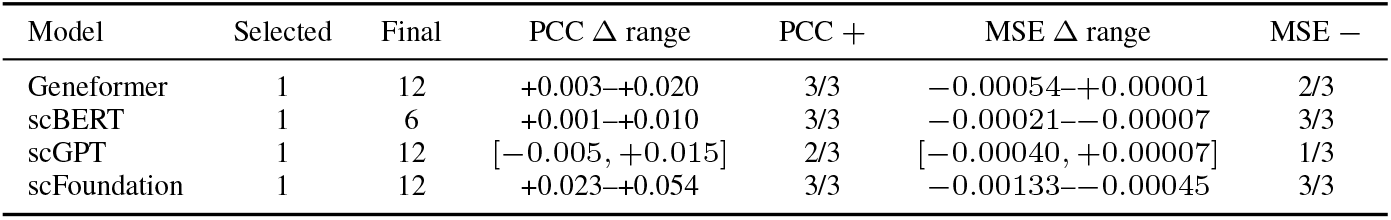
Perturbation LODO results across three held-out datasets. Entries show the range of selected-minus-final differences across held-out datasets.

Across the 12 primary perturbation model–dataset settings, at least one non-final layer improved a retrieval metric in 11 settings, and no evaluated layer triggered a collapse warning. The warning rule was applied to predicted-to-observed variance ratio, mean pairwise Euclidean-distance ratio, and unique Pearson top-1 assignment fraction. These diagnostics support the interpretation that early-layer performance was not explained solely by mode collapse, while not establishing universal early-layer superiority.

**Appendix Figure 1:**
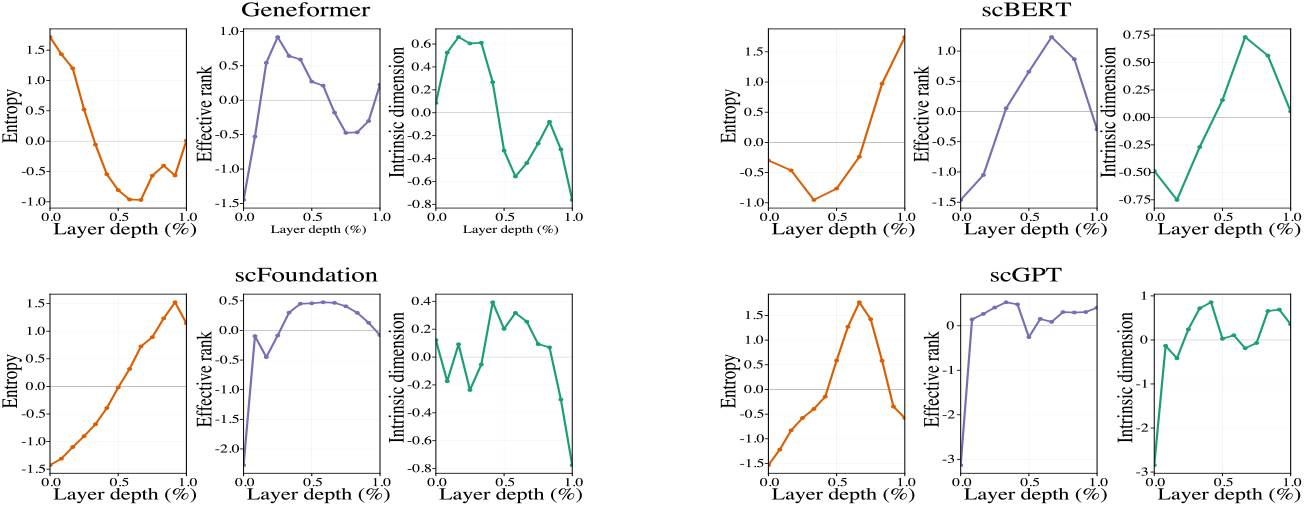
Layer-wise representation diagnostics for cell type prediction, averaged across datasets for each scFM. The panels show entropy, effective rank, and intrinsic dimension profiles across normalized depth.

**Appendix Figure 2:**
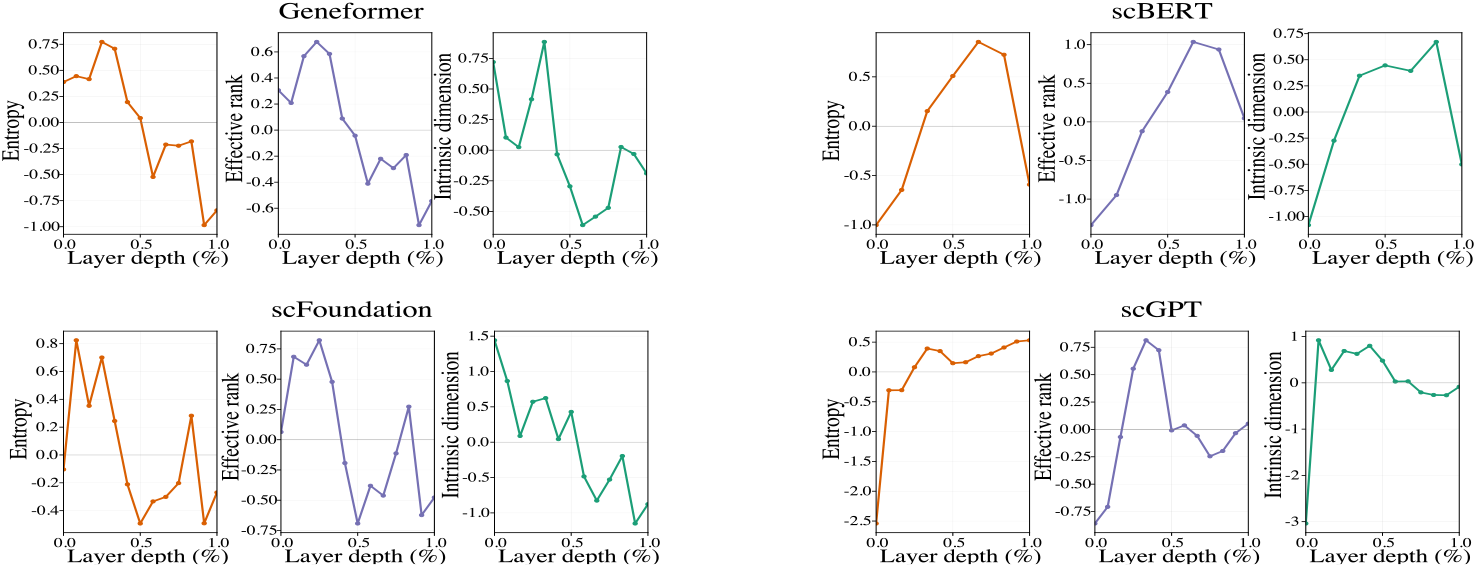
Layer-wise representation diagnostics for perturbation-response prediction, averaged across datasets for each scFM. The panels show entropy, effective rank, and intrinsic dimension profiles across normalized depth.

**Appendix Figure 3.**
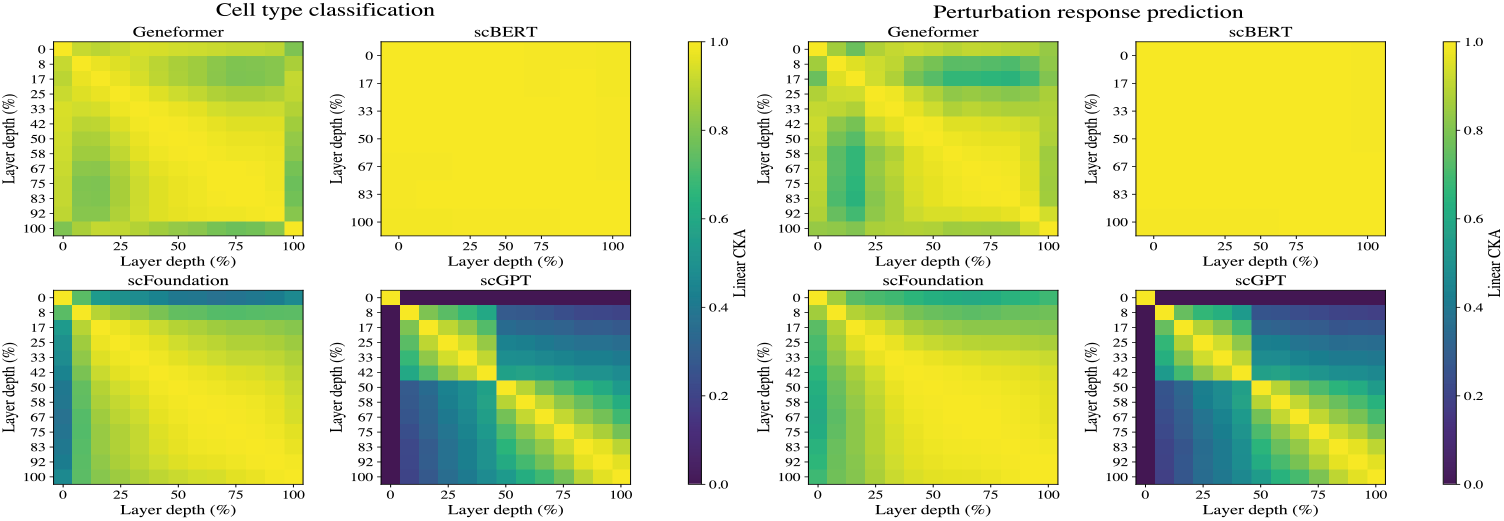
Average layer-wise CKA heatmaps across datasets for the four scFMs. The left panel summarizes cell type prediction datasets and the right panel summarizes perturbation-response datasets.

## Notes

### Competing Interest Statement

The authors have declared no competing interest.

